# Wasted efforts impair random search efficiency and reduce choosiness in mate-pairing termites

**DOI:** 10.1101/2024.02.01.578198

**Authors:** Nobuaki Mizumoto, Naohisa Nagaya, Ryusuke Fujisawa

## Abstract

Random search theories predict that animals employ movement patterns that optimize encounter rates with target resources. However, animals are not always able to achieve the best search strategy. Energy depletion, for example, limits searchers’ movement activities, forcing them to adjust their behaviors before and after encounters. Here, we quantified the cost of mate search in a termite, *Reticulitermes speratus*, and revealed that the searching cost reduces the selectivity of mating partners. After a dispersal flight, termites search for a mating partner with limited reserved energy. We found that their movement activity and diffusiveness progressively declined over extended mate search. Our data-based simulations qualitatively confirmed that the reduced movement diffusiveness decreased the searching efficiency. Also, prolonged search periods reduced colony foundation success and the number of offspring. Thus, mate search imposes doubled costs on termites. Finally, we found that termites with an extended mate search reduced the selectivity of mating partners, where males immediately paired with any encountering females. Thus, termites dramatically changed their mate search behavior depending on their physiological conditions. Our finding highlights that accounting for the searchers’ internal states is essential to fill the gap between random search theories and empirical behavioral observations.

## Introduction

When searching for targets whose location is unknown, animals benefit by adopting movement patterns that promote random encounters (Bartumeus and Catalan, 2009; Viswanathan et al., 2011). Random search theories have predicted the best movement patterns that maximize encounter rates with targets across various environmental conditions, including distribution and movement patterns of targets (Bartumeus et al., 2005; Cain, 1985; Palyulin et al., 2014; Viswanathan et al., 1999) and ecological contexts (Abe, 2020; Abe and Shimada, 2015; Fagan et al., 2020; Mizumoto et al., 2017a). Also, empirical studies have shown that animals use optimal random search strategies by focusing on how searchers change their movement patterns across differential searching environments (Bartumeus et al., 2003; Humphries et al., 2010; Mizumoto and Dobata, 2019; Reijers et al., 2021; Weimerskirch et al., 2007). However, although these studies emphasized the importance of external environments to determine the optimal ransom movement patterns, the internal conditions of searchers have rarely been considered (Joo et al., 2022). Real animals have different entities, such as sexes and ages, among individuals, and their physiological conditions dramatically change according to time due to, e.g., changes in nutritional conditions. Such biological contexts are essential to connect theories of random search and empirical observations of animal searching behaviors.

Mate search is essential for sexual reproduction, and random search theories predict that both females and males evolved sex-specific movement patterns to optimize mating encounters (Mizumoto et al., 2017a; Mizumoto and Dobata, 2018; Reynolds, 2006). However, behavior after encounters is also critical for a successful mating. When mating costs are high, both females and males decide whether to accept or reject the partner after encountering each other (Courtiol et al., 2016). This choosiness can incur a fitness cost even in monogamous animals because of the sequential mating encounters (e.g., (Dechaume-Moncharmont et al., 2016), but see (Forstmeier et al., 2021)). Mate searchers may not revisit rejected partners; thus, rejecting mates increases the probability of dying before being reproduced (Etienne et al., 2014; Kokko and Mappes, 2005). The level of choosiness is highly variable according to species (e.g., (Edward and Chapman, 2011)), and theory predicts that it depends on a variety of factors, such as the frequency of mating encounters (Courtiol et al., 2016; Etienne et al., 2014). Thus, by changing the encounter frequency, random search efficiency must subsequently affect the decision-making process for mate choice. For example, if mate searchers could not achieve the optimal movements due to physiological conditions, mate searchers would plastically adjust their subsequent mate choice after encounters.

To study the plastic changes of random search, *Reticulitermes* termites provide an ideal system. During a particular season, alates (winged reproductives) fly to disperse for mating. After dispersal, termites shed their wings, walk to search for mating partners, and, upon successful encounters, perform tandem runs to stay together while looking for a potential nest site (Fig. 1A) (Nutting, 1969). In this process, both females and males of termites flexibly change their random search strategies to enhance mating encounters. Before encounters, both females and males actively explore the environments to cover a broad area (Mizumoto and Dobata, 2019), while leaders pause and followers move to enhance reunion once they accidentally separated during tandem pairing (Mizumoto et al., 2022; Mizumoto and Dobata, 2019). Pairs excavate wood or soil to establish a nest when they find a candidate place. Because termites are monogamous and spend the rest of their life with a specific partner, females and males are expected to choose a partner to some extent (Li et al., 2013; Matsuura et al., 2002). However, at the same time, as their mate search period is strictly limited from a few hours to a few days (Kusaka and Matsuura, 2017), choosy individuals can likely end up having no partner and die. Thus, termites are expected to change their choosiness dynamically, depending on the availability of mating partners surrounding themselves. A previous study consistently shows that the random search strategy of termite males depends on the local density of females (Mizumoto et al., 2020). Note that males, rather than females, actively choose partners in termite species with female-led tandems. Tandem runs do not start without males following females (Mizumoto et al., 2022), and females do not change their behavior according to their partners (Mizumoto et al., 2021, 2020).

**Figure 1.**
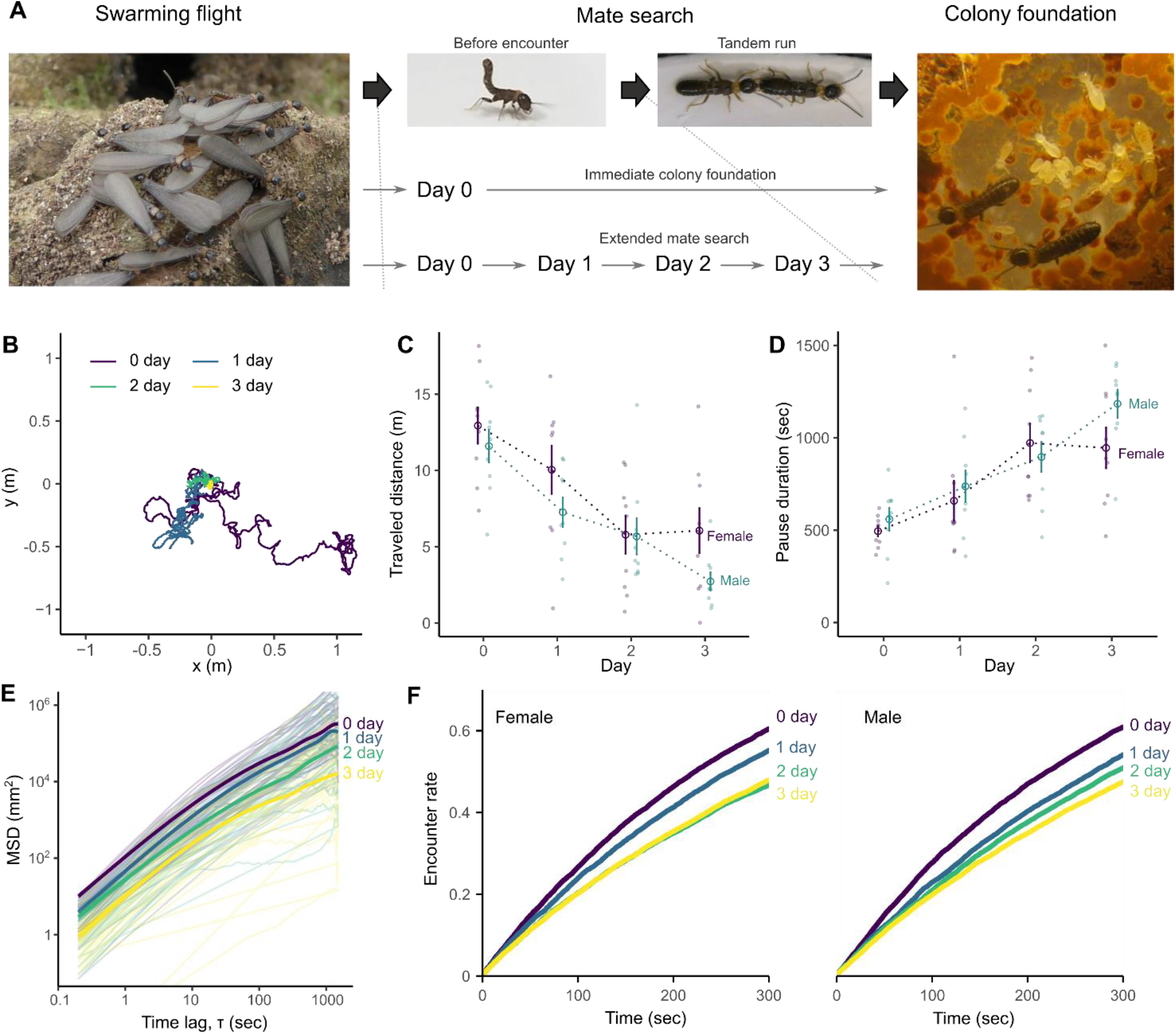
Change in movement patterns according to extended mate search in a termite *Reticulitermes speratus*. (A) Overview of the mating biology and experimental setup. After swarming flights, termites shed their wings and walk to search for a partner. After a successful encounter, they formed a tandem running pair and found a colony. Mate search usually ends in a day; otherwise, it can last until they find a partner. We observed their movement patterns every 24 hours for 30 minutes over four days (day 0-3). (B) Representative trajectories of a termite. Each trajectory corresponds to 25 minutes, where a termite moved less and less distances across days. (C-D) Comparison of traveled distances (C) and pausing duration (D) of termites across days after swarming. Bars indicate mean ± s.e. (E) Mean squared displacements (MSD) of the trajectories on the servosphere across days after swarming. Thick lines indicate the mean for each day’s observations, where data of females and males were pooled. (F) Simulation results for comparing searching efficiencies of movement patterns observed in termites across days after swarming (*L* = 223.6, φ = 7). The results were obtained from the means of 100,000 simulations for each combination.

In this study, we quantify the cost of mate search and investigate its influence on the mate-searching behavior in a termite, *Reticulitermes speratus* (Kolbe 1985). Mate search is the only behavior of the termites dealates, and mate search will last until they find a partner; otherwise, they die (Mizumoto et al., 2016). If they could not find a partner, the mate search could last multiple days with photoperiodic circadian rhythms (Mizumoto et al., 2017b). Thus, we forced dealates to do mate searches without encountering partners for 72 hours and examined how this extended search alters their movement patterns, colony foundation success, and mate choice. The mate search could be highly costly, given that termites do not feed to obtain energy until they start a new colony and have workers for foraging (Chouvenc, 2022; Inagaki et al., 2020), and social isolation is exceptional in their life history (Koto et al., 2015). Thus, the costs associated with mate search must strongly impact their searching and mating strategies.

## Results

### Change of searching activity over time

After the swarming flights, both females and males actively move to search for a mating partner in *R. speratus* (Mizumoto and Dobata, 2019). However, according to time, the search activity was progressively reduced (Fig. 1B). When we observed mate search behaviors of termites on a servosphere each day, we found that traveled distances significantly declined according to extended mate search (LMM, LRT, day: χ^2^_3_ = 98.9, *P* < 0.001, Fig. 1C), with females moving more distances than males (χ^2^_1_ = 4.5, *P* = 0.033). Similarly, both females and males paused for a longer duration according to observational day (LMM, LRT, day: χ^2^_3_ = 71.7, *P* < 0.001, Fig. 1D), with no sexual differences (χ^2^_1_ = 1.84, *P* = 0.17, Fig. 1D). This difference of movement activities turned out the different diffusive properties of termites. Termites just after swarming had the largest mean squared displacements (MSD), and the value of MSD decreased according to days after swarming (LMM, LRT, day: χ^2^_3_ = 398.9, *P* < 0.001, Fig. 1E). Note that the slopes did not change according to observation days in termites (LMM, LRT, interactions between day and τ: χ^2^_3_ = 5.89, *P* = 0.11), indicating that the decrease of MSD came from the changes of movement activities, such as movement speed and pausing duration, rather than the type of random walks, e.g., changes of turning patterns or Lévy walk properties (Viswanathan et al., 2011). Therefore, the observed behavioral changes did not reflect the changes in random search strategies but the inactivity of termites, perhaps caused by energy depletion.

When termites search for a mating partner whose location is unknown to searchers, high diffusiveness (large MSD) is critical for encounter efficiency (Mizumoto and Dobata, 2019), as shown in theoretical studies on random search when searchers do not have any prior information on the location of randomly distributed targets (James et al., 2008; Mizumoto et al., 2017a). Because termites moved fewer distances with less diffusive properties with longer pauses (Fig. 1B-E), searching efficiency is expected to reduce according to extended mate search. Accordingly, our data-based simulations (see Methods) demonstrated that searching efficiency of termites was progressively reduced according to observation days (Fig. 1F). In both females and males, searching efficiency was highest in termites just after the swarming, which showed the highest diffusive properties (Fig. 1EF), and lowest in termites after three days that had the lowest diffusive properties (Fig. 1EF).

#### Fitness cost of extended mate starch

If the inactivity of termites with extended mate search comes from energy depletion, extended search causes long-term fitness costs, in addition to the reduction of searching efficiency. We found that extended mating search incurs the cost of colony foundation success. The colony foundation success was significantly reduced in termite pairs that experienced three days extended search, compared with those found a colony just after swarming (GLMM, LRT, sex: χ^2^_1_ = 4.02, *P* = 0.04; Fig. 2A). Also, after the successful colony foundations, pairs with extended mate search had the significantly smaller number of offspring (GLMM, LRT, sex: χ^2^_1_ = 4.14, *P* = 0.04, Fig. 2B). Thus, even after the mate search period, extended mate search adversely impacted the performance of the colony foundation.

**Figure 2.**
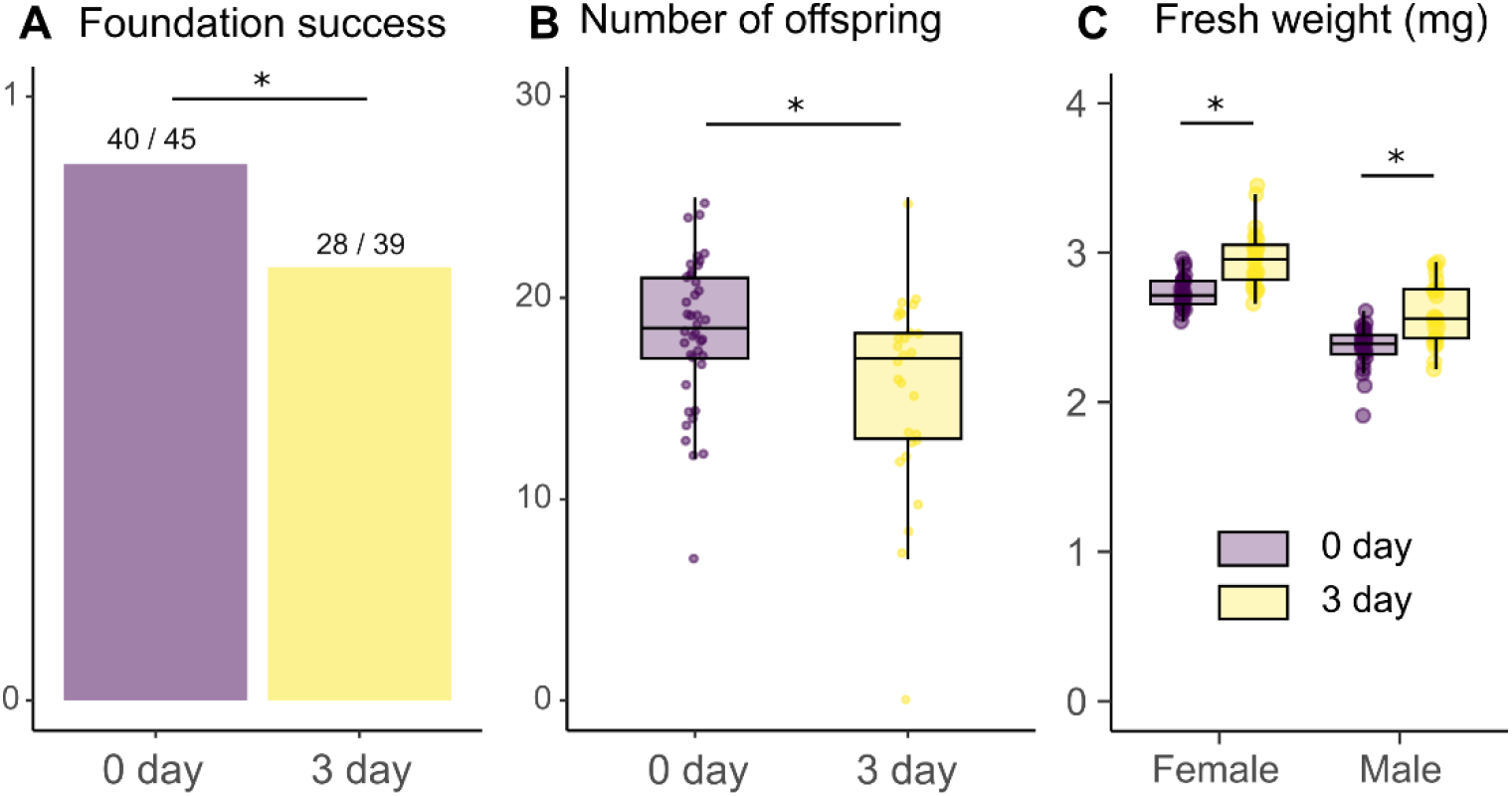
The long-term cost of extended mate search in termites. (A) Comparison of colony foundation success and (B) number of offspring between alates just after swarming (0-day) and after three days of mate search (3-day). (C) Comparison of fresh body weight between alates just after swarming (0-day) and after three days of mate search (3-day). * indicates statistically significant differences (GLMM or LMM, *P* < 0.05).

The termites with extended mate search should be less attractive options for other termites to make a pair because they perform less during colony foundations (Fig. 2AB). We then tested if the extended search incurred the additional cost by reducing the competitiveness during partner selection. We presented two females (or two males) who experienced extended mate search (old) or just after swarming (new) to a new male (or a female). Then, we observed which tandem pairing, new-new or new-old, was observed after 10 minutes. We found that termites with extended mate search performed as well as the termites just after swarming as a tandem pairing partner candidate during mate choice. There was no significant difference between old and new males (Binomial test, old:new = 19:10, *P* = 0.14) and between old and new females (Binomial test, old:new = 17:12, *P* = 0.46).

Interestingly, termites after extended mate search had larger fresh body weight than termites just after swarming (LMM, LRT, sex: χ^2^_1_ = 153.12, *P* < 0.001, day: χ^2^_1_ = 67.04, *P* < 0.001, interaction: χ^2^_1_ = 0.02, *P* = 0.88, Fig. 2C). Termite alates lose weight by evacuating water just before the swarming to improve their flight and dispersal capacity. Thus, termites with extended mate search could gain weight by getting water in their body (Chouvenc, 2022). Because heavier termites often tend to obtain a partner if they have multiple individuals around (Li et al., 2013; Matsuura et al., 2002), termites with extended mate search might compensate for the cost of mate search by collecting water.

#### Changes in tandem pairing after extended mate search

Finally, we investigated how extended mate search alters the decision-making of termites after encountering mating partners. We introduced 40 termites (20 females and 20 males) that experienced extended mate search or just after swarming to a large experimental arena (ø = 600 mm) and then observed the dynamics of tandem pairing. We found that extended mate search in isolation dramatically increased the motivation of tandem running behavior in termites (Video S1-2). In total, we observed 219 tandem runs for termites just after swarming (166 heterosexual, 36 male-male, one female-female tandem run, and 16 tandems with >2 individuals), while 679 for termites that experienced 72 hours of mate search (564 heterosexual, 30 male-male, six female-female, and 79 tandems with >2 individuals), during 30 minutes observations. Termites with extended mate search showed significantly more heterotrophic tandem runs (Wilcoxon Signed rank test, *V* = 0, *P* = 0.03; Fig. 3A) and tandem runs with more than two individuals (Wilcoxon signed rank test, *V* = 0, *P* = 0.03; Fig. 3A), but not male-male (Wilcoxon signed rank test, *V* = 14, *P* = 0.56; Fig. 3A) and female-female tandem runs (Wilcoxon signed rank test, *V* = 1.5, *P* = 0.375; Fig. 3A). Most tandem runs with more than two individuals were that two males following one female (84/95; 88.4%). There was no significant difference in the number of tandems between nest-mate or non-nest mate pairing in all these pairing combinations (Wilcoxon signed rank test, 0 day: *V* = 10, *P* = 1; 3 day: *V* = *8, P* = 0.6875).

**Figure 3.**
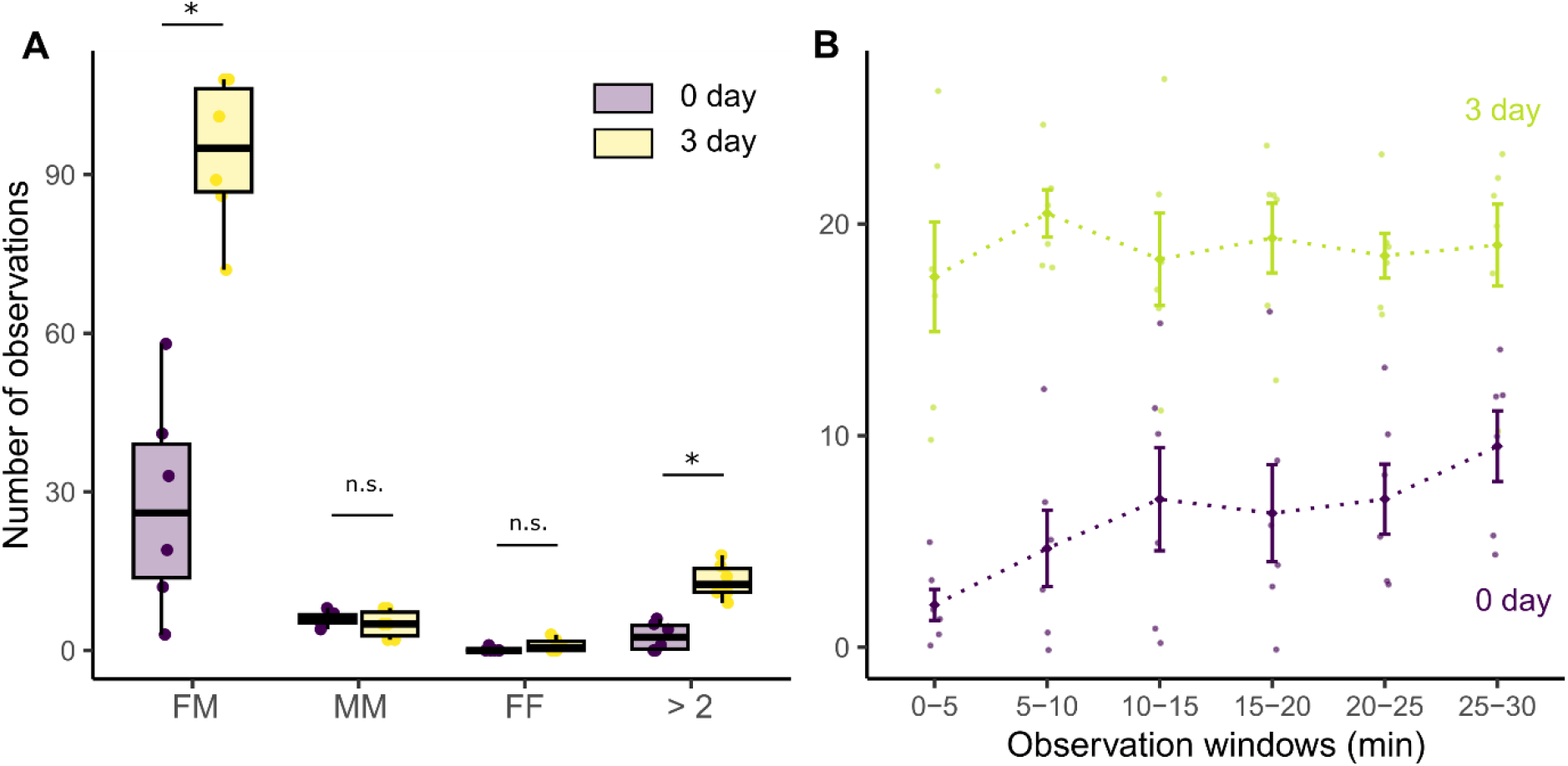
Comparison of observed tandem runs between termites just after swarming (0-day) and termites that experienced isolated mate search for 72 hours (3-day). (A) Comparison of the total number of observations for each unit during 30 minute observations. FM: female-male, MM: male-male, FF: female-female, and >2: runs involving more than two individuals. “*” indicates statistical significance (Wilcoxon signed rank test, *P* < 0.05), while “n.s.” indicates non-significance (> 0.05). (B) Time development of the observed number of tandem runs. Error bars indicate mean ± S.E.

Furthermore, we found distinct patterns of time development of tandem pairing between termites with and without extended mate search. In termites just after swarming, the number of observed tandem runs increased according to the observational periods, where termites exhibit fewer tandem runs soon after being introduced to the arena (Spearman’s rank correlation, *S* = 3.55, *P* = 0.01; Fig. 3B). On the other hand, in termites with extended mate search, the number of tandem runs was higher than the last 5 minutes of termites just after swarming and did not increase according to observational periods (Spearman’s rank correlation, *S* = 26, *P* = 0.66; Fig. 3B). Thus, the motivation of tandem runs was highest in termites with experiencing extended searches, while it increased according to time in termites just after swarming.

## Discussion

Mate search incurs the doubled costs to termites. First, termites traveled less distance and discounted the diffusive properties after the extended mate search period (Fig. 1). As fast and diffusive movements are beneficial to increase encounter efficiency in termite random mate search (Mizumoto et al., 2020; Mizumoto and Dobata, 2019), reduction of movement capacity decreases mating encounters (Fig. 1F). Such inactivity of termites with extended mate search could be caused by the energy depletion (Wickman and Jansson, 1997) because termite reproductives use reserved fat body for dispersal flight and colony foundations and do not obtain further energy until they successful start establishing the new colonies (Chouvenc, 2022; Inagaki et al., 2020). Accordingly, energy costs associated with mate search further decreased the successes of colony foundations and the number of offspring after foundations (Fig. 2). Thus, mate search is a highly cost-intensive work for termites. Our results indicate that the search activity itself could lead to changes in movement patterns for a random search. Such time development of searchers’ internal state has rarely been considered in the theory of optimal random search. However, it could be a critical component in the movement ecology of real animals.

To compensate for reducing the probability of mating encounters due to inactivity, termites adjusted their mating preferences after the extended searches. Termites that experienced the extended searches had a much higher motivation for pairing than those just after swarming, where males followed any females upon encounters (Fig. 3, Video S2). This observation is consistent with the theoretical predictions that low choosiness is favored by a low encounter rate (Courtiol et al., 2016; Crowley et al., 1991; Etienne et al., 2014; Kokko and Johnstone, 2002; Real, 1990). Furthermore, in our observations of termites with extended mate search, two males frequently compete to obtain a single female (Fig. 3A, comparison of >2 observations, Video S2). Similar changes in mate choice preferences can be observed in other animals, e.g., the motivation of mating increases when the life expectancy is short (Wilgers and Hebets, 2012; Wilson et al., 2010), the cost of search is high (Alatalo et al., 1988; Milinski and Bakker, 1992; Willis et al., 2011; Wong and Jennions, 2003), and mating season is short (Gotthard et al., 1999). Furthermore, males isolated for long periods become less discriminative to their mating partners and often show same-sex pairing (Engel et al., 2015). Thus, termites do not solely rely on the random search strategy during mate pairing; instead, they combine it with other decision-making systems to rationally obtain a partner.

The pairing dynamics of termites were distinct between individuals just after swarming and with extended mate search. In termites just after swarming, the number of observed tandem runs increased during observational periods, while it was highest since the beginning in the individuals with extended mate search (Fig. 3B). This pattern could be interpreted with a sequential choice model (Janetos, 1980; Real, 1990). The sequential choice model is also called the “secretary problem,” where a decision-maker seeks to select the best option from a sequentially presented set of options. Like termite tandem pairing, the decision-makers must decide on encountering the option and cannot revisit the previous option once they reject it. Here, one of the optimal strategies is rejecting the first several options to learn the qualities of options and then making a choice based on the previous experience (Priklopil et al., 2015). In our experimental conditions, the density of termites is high, and males are expected to encounter many females just after swarming. Thus, they can evaluate multiple females before selecting a partner for tandem running. On the other hand, for males with extended mate search, the number of females they expect to encounter will be small as they already learned the density is low during isolation. Thus, males do not reject the first several partners for evaluation but accept any partners for tandem runs as soon as they encounter them. Our results shed light on the cognitive capacity of termite mate searchers.

In the theory of optimal random search strategies, conditions of individual searchers have rarely been taken into account, and searchers are supposed to exhibit the best performance during searching periods. However, our study showed that searching behavior itself can affect their movement capacity, altering their movement patterns and sequential decision-making after encounters. By filling the gap between theoretical studies on ideal searching conditions and empirical observations on realistic searching situations, our study contributes to understanding the evolution of random search strategies adapted in animals.

## Material and Methods

### Termite

We collected *R. speratus* alates with a piece of nesting wood from seven colonies in Kyoto in May 2016 and 2017. May is the swarming season of this species in Kyoto. All nesting wood pieces were maintained at 20°C until experiments. Before each experiment, we transferred nests to a room at 25°C, which promoted alates to emerge and fly. Alates were then collected and separated by sex. We used individuals who shed their wings by themselves. We stayed all experiments within 24 hours after the swarming flight.

### Change of searching activity over time

We investigated how searching activities change according to time after swarming flight using three colonies collected in 2016. We investigated the search activity of nine females and nine males (three from three colonies). Each individual was placed on an omnidirectional servosphere (Nagaya et al., 2017) and freely walked on an infinite two-dimensional surface for 30 minutes. The observation was performed four times every 24 hours, i.e., just after swarming (= day 0), day 1, day 2, and day 3. Termites were individually maintained in a Petri dish (φ = 90 mm) with a moistened piece of filter paper (a quarter of 70 mm). The bottom of the dish was polished so that termites could walk smoothly on it. We maintained each termite under the light condition of 16L8D and at 25°C during each observation interval. The observations were performed during the time of light conditions. Because data acquisition was not in a constant sampling rate in our servosphere, we interpolated the coordinates data to obtain the coordinates every 0.2 seconds (5Hz) after smoothing them with a median filter (k =5). Then. We removed data for the first five minutes and used the rest for 25 minutes for further analysis.

First, we obtained the moved distance every 0.2 seconds (steplength). We calculated the total distances termites walked during 25 minutes by summing up these steplengths. Also, we examined pausing duration during this period. In *Reticulitermes speratus*, the previous study estimated that the threshold for moving and pausing was 0.7mm (Mizumoto and Dobata, 2019); we regarded termites are moving if they moved more than 0.7mm in 0.2 seconds, while they are pausing if less than 0.7mm. We measured the total pausing duration during 25-minute observations. Then, we investigated the effect of time after swarming on walking distracts and pausing durations, using linear mixed models (LLM) that include time after swarming (0, 1, 2, and 3 days after swarming), sex, and their interactions as fixed effects, and the original colony as the random effect (random intercept). The statistical significance of each variable was tested using the chi-square test (type II ANOVA) herein and the following statistical analysis.

Second, we evaluated the diffusive properties of individual movements. High diffusiveness is critical for the efficiency of the random search when searchers do not have any prior information on the location of targets that are randomly distributed (James et al., 2008; Mizumoto et al., 2017a), which is relevant for termite mate search before encounters (Fig. 1A) (Mizumoto and Dobata, 2019). We computed the MSD to compare the diffusive properties of individual movements across time after swarming. The MSD is the mean of the squared distance that an organism travels from its starting location to another point during a given time, τ. We obtained MSD in the range of 0.2 < τ < 1500, using the function *computeMSD*() in the package “flowcatchR”. To compare the MSD between time after swarming, we used an LMM, where τ, time after swarming, sex, and their interactions were included as fixed effects, and individual IDs nested within original colonies were included as a random effect (random intercept). MSD and τ were log transformed before the LMM fitting.

### Evaluation of searching efficiency

Because termites moved fewer distances with less diffusive properties and many pauses (Fig. 1B-E), searching efficiency is expected to reduce according to extended mate search. To quantify the search efficiency, we used a data-based simulation approach. After randomization, we projected the trajectories of a female and a male and calculated if and when they encountered. When mate search of termites extended to multiple days, they synchronized their search efforts with the following swarming events (Mizumoto et al., 2017b). Given that most termites do not extend mate search for multiple days due to pairing or predation (Mizumoto et al., 2016), we expect that mating encounters of extended mate searchers are usually with new mate searchers (day 0 individuals). Thus, we investigated the encounter efficiency in the combinations of 0 day-0 day, 0 day-1 day, 0 day-2 day, and 0 day-3 day trajectories. We picked up one trajectory with 5 FPS of a female and a male and placed them randomly in a periodic boundary condition of size = *L* × *L*. Each trajectory was horizontally and vertically reversed with the probability of 1/2. Following inversion, we rotated the trajectory at random degrees from 0 to 360 around the starting point of the trajectory. After projecting two trajectories, we estimated if and when these two individuals encountered. We regarded they encountered when two are within the distance φ at the same time. We performed this process 100,000 times for eight cominations (0 day-0 day, 0 day-1 day, 0 day-2 day, and 0 day-3 day, for female-male and male-female). The parameter φ value was set as 7 mm, based on the body size of *R. speratus* (Mizumoto and Dobata, 2019), and *L* as 223.6, based on the previous studies (Kusaka and Matsuura, 2017; Mizumoto and Dobata, 2019).

### Fitness cost of extended mate starch

We investigated the long-term fitness cost of extended mate search after colony foundation using four colonies collected in 2017. We prepared termite females and males that were isolated for 72 hours in a Petri dish described above. After 72 hours of isolated mate search, these termites were paired with each other. Similarly, we also prepared pairs of termites just after swarming. Each pair was introduced to a Petri dish (φ = 40 mm) filled with brown-rotted pinewood mixed cellulose medium (Mitaka et al., 2023) at a depth of 5 mm, where termites excavated into the medium to establish their nests. All pairs were produced using nest-mate individuals. We prepared 12 pairs for each condition in three colonies and nine pairs for one colony. As several individuals were dead during 72 hours of isolated mate search, we had 45 pairs just after swarming and 39 pairs after 72 hours of matter search in total.

All dishes were maintained at 25? in a dark condition for 60 days. After 60 days, we opened all dishes and counted the number of surviving individuals (female, male, larvae, and eggs, separately). We defined that the pair succeeded in colony foundation only when both the female and the male survived. We compared colony foundation success between just after swarming and 72 hours after swarming, using a generalized linear mixed model (GLMM) with binomial distribution and logic link function, in which termite condition was included as a fixed effect, while the original colony was included as a random effect (random intercept). We also compared the number of eggs and larvae between termite conditions among pairs that succeeded in colony foundation. We used a GLMM with Poisson distribution; otherwise, it was the same as the other one.

### Change of fresh weight and mate preference

Fresh body weight is often used as an indicator of the quality of termite dealates because heavier termites are preferred as partners (Li et al., 2013; Matsuura et al., 2002). We examined how body weight could change during extended mate search and how it affected mating preferences. Using three colonies collected in 2016, we compared the fresh weights of eight females and males for each colony just after swarming and after 72 hours of mate search. We performed the measurements on a scale of 0.01 mg.

To investigate the mate preference of termites, we observed the mate competition situations between termites just after swarming (new) and after 72 hours of mate search (old). We introduced three termite dealates (new female + old female + new male, or new female + new male + old male) in a petri dish arena (φ = 90 mm) filled with moistened plaster. After 10 minutes of introduction, we investigated the pairing combination. All individuals were painted and marked with one color on the abdomen for individual identification (PX-20; Mitsubishi). We used a binomial test to test the bias in pairing combinations (old or new). All three individuals were from different colonies.

### The effect of extended searching on tandem pairing occurrence

We investigated how extended mate search affects the pairing dynamics, using colonies collected in 2016. After swarming, ten females and ten males of dealate were selected from two different colonies. These 40 individuals were painted and marked with one color on the abdomen for sex and colony identification (PX-20; Mitsubishi). All 40 individuals were maintained for 30 min (just after swarming) or 72 hours (extended mate search) separately in a 24-well plate before the observation. Note that the results of the 30-minute treatment were reported in a previous study (see Text S1 of (Mizumoto et al., 2022)). We introduced 40 individuals to the experimental arena (ø = 600 mm). The experimental arena consisted of a Styrofoam board (600 × 600 mm) and a circular plastic tube (ø = 600 mm, height = 100 mm). After being introduced in the arena, we observed termite movements for 30 minutes within a part of the experimental arena (200 × 100 mm) located at the edge of the circular arena (Fig. S2). We did so because most individuals walked along the arena’s edge, repeatedly passing across the observational arena (Video S1 and S2). We counted the number of individuals passing across the observational area with their status (single individuals, heterosexual tandems, same-sex tandems, tandems with >3 individuals). We performed the experiments six times with different colony combinations for each treatment. The experimental arena was cleaned with 70% ethanol and distilled water before each experiment.

We used the Wilcoxon signed rank test to compare the number of observations of pairing units between termites just after swarming and after extended mate search. We also investigated the time development of observed tandem running pairs. We binned our 30-minute observation into 0-5 minutes, 5-10 minutes, …, 25-30 minutes, and counted the number of observed tandem pairs during each time-windows. We used Spearman’s rank relation test to test if the number of observed tandem running pairs increased according to time developments.

All analyses were performed by R v4.3.1 (R Core Team, 2023) with libraries of “lme4,” “car,” and “exactRankTests” for statistical tests and “Rcpp” for data-based simulations. All data with R and Cpp scripts are available at GitHub (github.com/nobuaki-mzmt/termite-mate-search-cost). The accepted version will be uploaded to Zenode to obtain DOI for the version of the record.

## Acknowledgments

We thank Shigeto Dobata, Kenji Matsuura, and Tomonari Nozaki for their valuable intellectual feedback on the experiments and Tadahide Fujita and Tomonari Nozaki for their help with termite collections. We also appreciate Seagate Recovery Services (https://www.seagate.com/products/rescue-data-recovery/) for recovering our data after an unexpected loss and emphasizing the importance of backups. This study was supported by Grants-in-Aid for JSPS Research Fellow 15J02767 (NM), Grant-in-Aid for Scientific Research (C) 23K03776 (RF), Grant-in-Aid for Scientific Research (B) 21H01295 (NN), and an IPSF fellowship from OIST (NM).

## Author Contributions

N.M.: conceptualization, data curation, formal analysis, funding acquisition, investigation, methodology, project administration, resources, supervision, validation, visualization, writing-original draft, writing-review and editing. N.N.: resources, software, writing-review and editing. R.F.: resources, writing-review and editing.

