## Supplementary material for "Wasted efforts impair random search efficiency and reduce choosiness in mate-pairing termites": Fig. S1-2

This file contains

- **Figure S1**. Trajectories on the servosphere of all observed individuals.
- **Figure S2.** Observational arena for tandem pairing.
- Legends for Supplementary Video S1 and S2

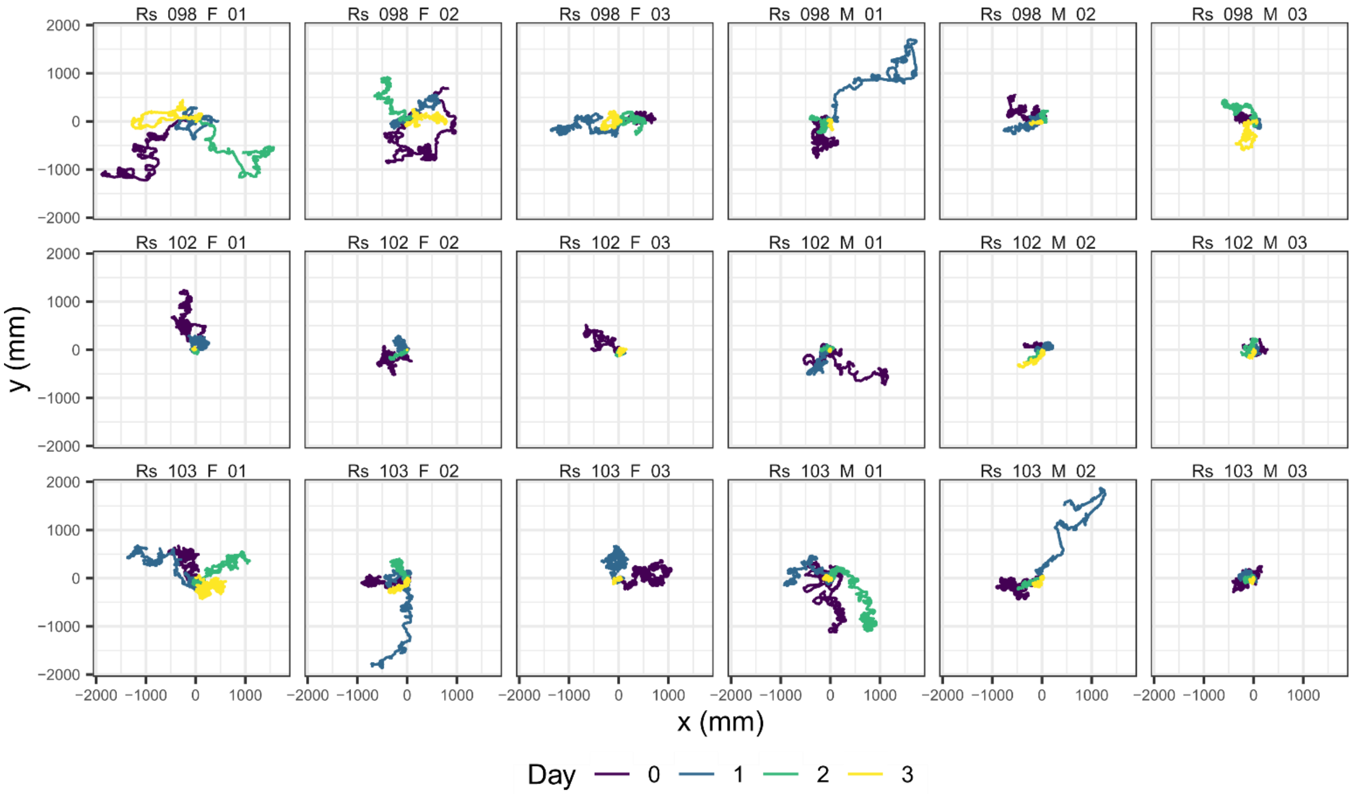


**Figure S1**. Trajectories on the servosphere of all observed individuals. Each label indicates Species name (Rs = *Reticulitermes speratus*) + Colony name + Sex + Replicates.

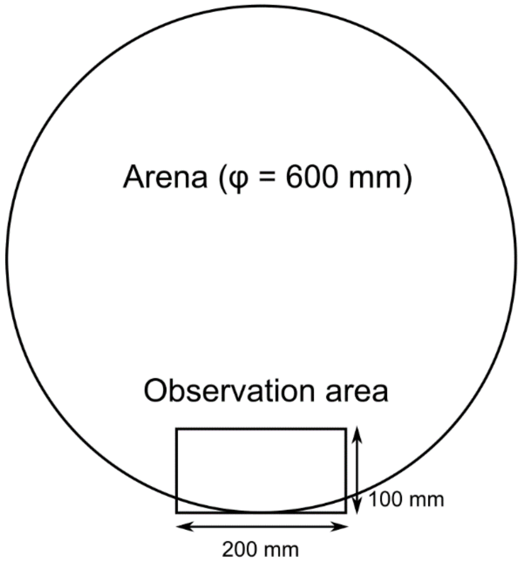


**Figure S2.** Observational arena for tandem pairing. A video camera was located above the observation area on a tripod. Termites were released at the center of the arena, and those who crossed the observation area were observed.

**Video S1.** The clip of the observation arena for termites just after swarming. Pink and yellow indicate females, while blue and green indicate males.

**Video S2.** The clip of the observation arena for termites with extended mate search. Pink and yellow indicate females, while blue and green indicate males.
